# proovframe: frameshift-correction for long-read (meta)genomics

**DOI:** 10.1101/2021.08.23.457338

**Authors:** Thomas Hackl, Florian Trigodet, A. Murat Eren, Steven J. Biller, John M. Eppley, Elaine Luo, Andrew Burger, Edward F. DeLong, Matthias G. Fischer

## Abstract

Long-read sequencing technologies hold big promises for the genomic analysis of complex samples such as microbial communities. Yet, despite improving accuracy, basic gene prediction on long-read data is still often impaired by frameshifts resulting from small indels. Consensus polishing using either complementary short reads or to a lesser extent the long reads themselves can mitigate this effect but requires universally high sequencing depth, which is difficult to achieve in complex samples where the majority of community members are rare. Here we present *proovframe*, a software implementing an alternative approach to overcome frameshift errors in long-read assemblies and raw long reads. We utilize protein-to-nucleotide alignments against reference databases to pinpoint indels in contigs or reads and correct them by deleting or inserting 1-2 bases, thereby conservatively restoring reading-frame fidelity in aligned regions. Using simulated and real-world benchmark data we show that proovframe performs comparably to short-read-based polishing on assembled data, works well with remote protein homologs, and can even be applied to raw reads directly. Together, our results demonstrate that protein-guided frameshift correction significantly improves the analyzability of long-read data both in combination with and as an alternative to common polishing strategies. Proovframe is available from https://github.com/thackl/proovframe.

## Introduction

The advent of long-read sequencing as a complementary approach to high-throughput short-read sequencing has played a key role in shaping the postgenomic era^1^. The two widely available long-read sequencing technologies from Pacific Biosciences and Oxford Nanopore Technologies now readily produce reads several tens of kilobases in length^2,3^, thereby providing the means to overcome long-standing assembly problems such as genomic repeats or structural heterogeneity^4–6^. This has opened new ways to study genomes across fields and scales -from viral outbreaks^7^ and antibiotic-resistant bacteria^8^ to eukaryotic genomes^9–11^, complex symbiotic partnerships^12,13^ and natural viral populations^14^.

However, the advantages of long reads come at a price. Despite recent advances in engineering and chemistry, the error rates for the technologies remain an issue limiting their usability: State-of-the-art PacBio data reaches accuracies of 85-90% with errors comprising around 2% substitutions, 8% insertions, and 3% deletions, while current Nanopore data achieves accuracies of 85-95% with around 3-5% substitutions, 2-4% insertions and 2-4% deletions^2,3^. Various approaches for pre- and post-assembly correction by consensus polishing have been devised to mitigate these errors, including hybrid correction with short reads or self-correction among overlapping long reads^15–20^. But even for assembled and polished data, frameshifts remain an issue particularly for assemblies generated from long-read data alone^21^. As an additional strategy, these frameshifts can be identified and corrected by comparison to reference protein databases, as implemented in the microbiome analysis platform MEGAN-LR^22^ using the frameshift-aware aligner DIAMOND^23^. All of these approaches, however, rely on sufficient coverage in either the long-read data itself or the accompanying short-read data for the computation of read-to-read alignments and the subsequent inference of consensus sequences with high accuracy.

In metagenomics projects, which can benefit significantly from long reads by way of improved binning and the recovery of high-quality genomes^24,25^, achieving sufficient coverage for assembly and consensus-polishing is not a given. In many natural communities, only a few species are abundant while the majority are rare^26^. Even among abundant members, high genetic complexity, such as observed for viral populations^14^, or localized intra-population heterogeneity tied to, for example, niche-differentiation^27^ or defense^28^, can render significant fractions of the data unpolishable.

Proovframe offers an accurate and comprehensive strategy to correct frameshift errors in protein-coding regions of long-read sequencing results by using distantly related protein sequences as guides. Since it does not require short-read sequencing data for polishing and can work with low-coverage long-reads, proovframe can yield substantial improvements for long-read sequencing of genomes and metagenomes at a low cost.

## Implementation

### Frameshift-aware read-to-protein alignment

Proovframe’s correction procedure consists of two steps: a frameshift-aware nucleotide-to-protein alignment and the subsequent alignment-guided correction of the erroneous nucleotide query sequences. Queries can be anything from raw reads to polished contigs. To compute frameshift-aware nucleotide-to-protein alignments we use the high-performance aligner DIAMOND^23^. DIAMOND produces BLAST-like local sequence alignments and provides different sensitivity modes that allow - at the expense of speed - alignments of sequences with as little as 30% sequence identity. Only the aligned regions of proteins are used to guide corrections, and hence for diverged reference proteins, correction of only parts of genes is possible.

Recent versions of DIAMOND (v0.9.14 and newer) provide a long-read mode that can model frameshifts in nucleotide-to-protein alignments (option ‘--frameshift 15’). For each query, we greedily collect only the best alignment for each location (option ‘--range-culling --max-target-seqs 1’) and retrieve the results in BLAST-like tab-separated format with an additional CIGAR-string column. The CIGAR string provides a compressed representation of the alignment, which in the case of DIAMOND also encodes the locations of the inferred frameshifts. This output file serves as input for the correction step.

With the ‘proovframe map’ command we provide a convenience wrapper to easily generate DIAMOND alignments with the appropriate parameters for a given set of input sequences and proteins. Although, if users prefer, alignments can also be computed with DIAMOND directly, which might provide additional control on, for example, distributing jobs to computing cluster infrastructures.

### Alignment-guided correction of reads

The correction of query sequences is carried out with the command ‘proovframe fix’, which takes as input the uncorrected query sequences and the nucleotide-to-protein alignments produced with DIAMOND, and returns corrected output sequences. For each query, all alignments are gathered and processed starting from the 3’ end of the query. Introducing modifications from the end of the sequence ensures that coordinates of loci not yet corrected remain stable. Using the alignment coordinates and the CIGAR string, we determine the location and nature (either a one- or two-base deletion or a one- or two-base insertion) of each frameshift-causing indel. We correct the indels by either the insertion of one or two “N” or the deletion of one or two bases at the given read location. For potentially larger indels, we also apply only one- or two-base modifications to restore frame fidelity. The rationale here is that we cannot robustly discriminate between the possibility of a larger insertion/deletion (which generally are rare) and the genuine lack/gain of an amino acid in the guide protein.

Generally, we found frameshift-locations determined by DIAMOND in the sequences to be fairly robust, although we note that different circumstances can introduce inaccuracies that affect correction: (a) If multiple indels occur within a few nucleotides, they can cancel each other out. For example, the net-change on frame-fidelity for a 1-base insertion next to a 1-base deletion is 0, and therefore, will not trigger a correction. The most likely outcome in this scenario is that the affected codons will encode for different amino acids, i.e. will introduce one, two, and in rare cases a few erroneous amino acids in the derived protein sequence (Fig. 1). (b) The exact location of deletions/insertions can be off by one or two bases due to ambiguities in the alignment. Hence, the modification of the sequence to restore the reading frame can in some cases introduce erroneous codons. (c) Frameshift locations can be off by one to a few codons (in rare cases) if the indel by chance resulted in codons that are consistent with the expected sequence at the respective locations. This effect is promoted by substitution errors present in the close vicinity of the frameshift causing sequences to align poorly in the first place. However, despite these issues, we found that the simple operations described above are sufficient to restore an overall high open-reading-frame quality.

**Figure 1.**
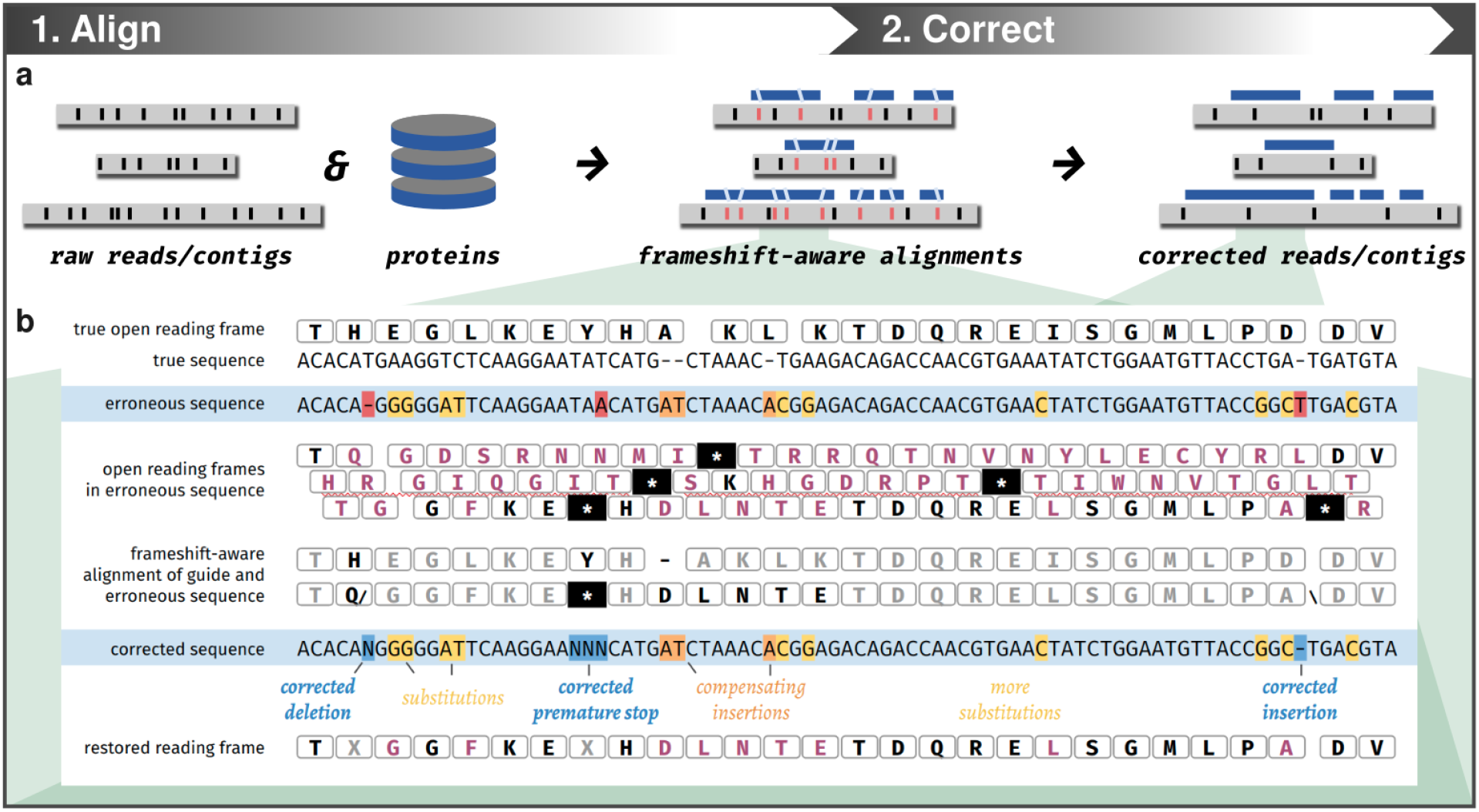
Proovframe’s workflow with an example of frameshift detection and reading frame correction. **a**) Overview of proovframe’s two-step correction procedure consisting of frameshift-aware nucleotide-to-protein alignments and the subsequent alignment-guided correction of query sequences (reads or contigs). Queries are shown as gray bars with errors as black ticks, guide-alignments as blue bars with frameshifts as light-blue slanted ticks. Red ticks indicate read errors identified as frameshifts from guide-alignment. **b**) Frameshift detection and correction for a short simulated sequence. Nucleotide sequences are shown as plain text, open reading frames as amino-acid translated codon sequences (boxed letters). The errorless nucleotide sequence and its translation are given at the top for comparison. Errors in the erroneous and corrected sequence are highlighted (red: correctable indel, orange: uncorrectable indel, yellow: substitution, blue: corrected indel or premature stop). Mismatched amino acids in open reading frames called on the raw and corrected sequence are shown in purple, premature stop codons as asterisks. Codons associated with frameshifts and premature stops are printed in black in the guide alignment.

In addition to insertion and deletion operations, we also scan the corrected sequences for the presence of internal stop codons, which we mask with NNN (Fig. 1b). These premature stops are in most cases either caused by substitution errors, complementing indels, or in some cases the frameshift corrections we performed. Masking them thus helps to restore the full open reading frame. Our approach, in general, is aimed at restoring frame-fidelity in the most conservative manner, i.e. introducing as few changes as possible, while still ensuring that the resulting nucleotide sequence encodes open reading frames that are in-frame with their guide proteins.

## Benchmark Datasets

To evaluate the performance of frameshift correction with respect to gene recall and gene call accuracy, we curated four benchmark datasets with different characteristics. The datasets together with reproducible code for their benchmarking are available from GitHub (thackl/proovframe-benchmark) and Zenodo (10.5281/zenodo.5164669)^29^.

### Viral genomes with simulated errors

For a set of eight published viral genomes^11^ we generated erroneous sequences with a nanopore-like error profile using NanoSim v2.6.0^30^. We called genes with Prodigal v2.6.3^31^ on the errorless reference and used the extracted proteins to correct the simulated reads with proovframe v0.9.5 running DIAMOND v2.0.4 for mapping. We then also called genes on the raw and corrected sequences and compared the results. False-positive gene predictions were defined as proteins called on the simulated and corrected sequences that could not be aligned back with DIAMOND to proteins from the reference. Gene predictions were analyzed in R and visualized with gggenomes^32^.

### *Mycoplasma bovis* reads corrected with UniRef90 proteins of different amino-acid identities

To test our method on real data, we downloaded recently published nanopore reads for *Mycoplasma bovis*^*33*^ (ENA study PRJEB38523, run ERR4179766). The reads were sequenced on an Oxford Nanopore MinION device and basecalled by the authors with bonito v0.1.3^34^. To assess the effect of protein divergence on frameshift correction, we used three different protein sets: the best matching proteins from the UniRef90 database^35^, as well as the best matching UniRef90 proteins with at most 80% or at most 60% identity to any protein from the reference genome. We obtained the latter sets by collecting all proteins from UniRef90 that align to the reference protein set, and then removing those above the identity threshold. Analogous to the viral test set, we called genes on an *M. bovis* raw and corrected read sets with Prodigal v2.6.3. For both nucleotide-to-protein alignments with DIAMOND and gene-to-protein translation, we used the non-standard genetic code for *M. bovis* (Genbank transl_table=4). For the visual comparison of gene predictions, we compared *M. bovis* reads longer than 15kbp to the respective locations on the reference genome. These locations were determined by mapping the reads to the reference genome with minimap2 v2.17 requiring a minimum alignment length of 90%.

### *Akkermansia* pangenomes

We extracted high-molecular-weight DNA from three fecal samples included in an Fecal Microbiota Transplantation (FMT) study by Watson et al^36^ using a protocol described by Trigodet and Lolans et al^37^. The three samples were collected from a single recipient (DA_R02 in the original study) before FMT (W0), one week after FMT (W1) and 48 weeks after FMT (W48). The Akkermansia population W1 that replaces W0 is also reconstructed from Illumina short reads and named as DA_MAG_00110 in the original study^36^. We used Flye v2.6 – which includes one round of long-read consensus polishing – for long-read assembly^20^, Pilon v1.23 for short-read polishing^38^, and proovframe v0.9.6 for frame-shift correction. To compute the pangenomes we used anvi’o v7^39,40^, which used Prodigal v2.6.3^31^ to identify open reading frames in each genome, DIAMOND v2.0.6^41^ to quantify the similarity between each pair of genes, the Markov Clustering Algorithm (MCL)^42^ to identify gene clusters, and muscle v3.8.1551 to align amino acid sequences^43^ to report a metric for ‘geometric homogeneity’, i.e., the proportion of gap characters in a given gene cluster (the URL https://merenlab.org/p serves a tutorial on anvi’o pangenomics workflow). We used the anvi’o program ‘anvi-display-pan’ to visualize pangenomes and finalized the figure for publication using Inkscape (https://inkscape.org/).

### Marine small-particle-fraction metagenomic reads

For this benchmark set we examined nanopore sequence data from a <0.2 µm small-particle marine sample collected from 25m at Station ALOHA (22° 45’ N 158° W) in the North Pacific ocean, fractionated as in Biller et al. 2014^44^, then extracted and sequenced as in Beaulaurier et al. 2020^14^. We randomly subsampled 1000 reads using seqkit v0.10.2^45^ and used a protein database that we assembled with the protein assembler PLASS sha-687d77^46^ from an accompanying Illumina HiSeq library made from the same DNA. We identified reads representing viruses with virsorter2 v2.1^47^ on both the raw and the corrected reads and annotated the reads flagged as viral with Prodigal v2.6.3^31^. The alignments we used for the correction, as well as the gene calls and virsorter classifications, are available from the repository.

## Evaluation

### Simulated viral nanopore sequences corrected with perfect proteins

To evaluate the performance of proovframe with respect to gene recall and gene call accuracy, we applied it to four test sets. First, we analyzed eight viral genomes^11^ with simulated nanopore-like errors. The observed error rates in the simulated data were 3.1% substitutions, 1.5% insertions, and 2.5% deletions. We called genes on the errorless reference and used the extracted proteins to correct the simulated reads with proovframe. We then also called genes on the raw and corrected sequences and compared the results (Fig. 2a/b).

**Figure 2.**
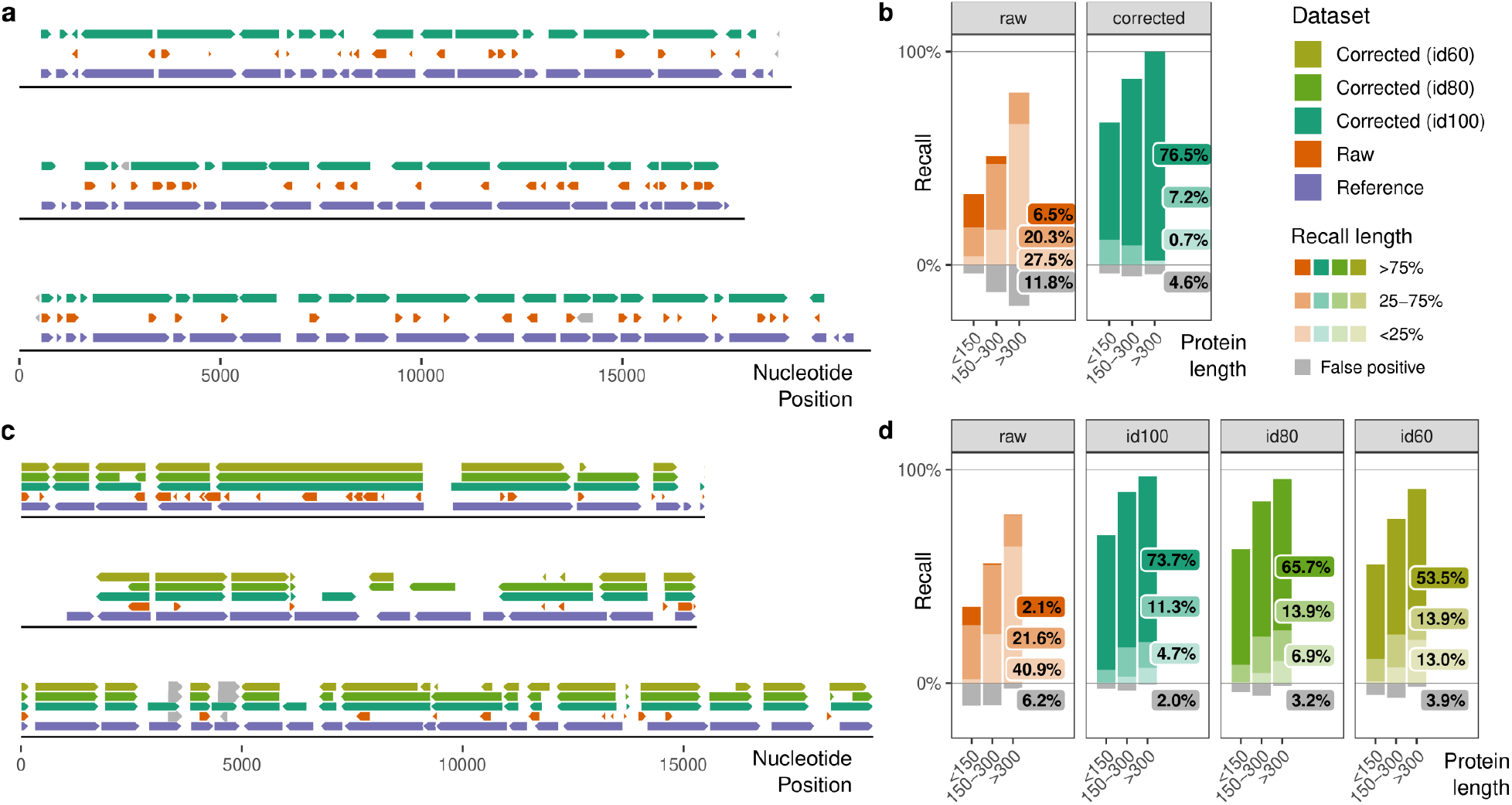
Frameshift correction results for simulated viral and real bacterial nanopore long reads. **a**) Comparison of gene predictions (arrows) made on three viral genomes containing no errors (purple), simulated nanopore errors (orange), and proovframe-corrected nanopore errors (green). Gray genes indicate false positive predictions, i.e. genes not found on the errorless reference. **b**) Histograms representing the fraction of genes of different length classes that were either predicted across less than 25%, 25-75%, and more than 75% of their expected length relative to the genes annotated on the reference. **c**) Comparison of gene predictions (arrows) made for three randomly chosen nanopore long reads of *M. bovis*, on a perfect reference (purple), and on real nanopore long reads before (orange) and after proovframe-correction with different protein sets: best-matching proteins from UniRef90 with at most 100%, 80% or 60% identity to any *M. bovis* reference proteins shown in teal, green and olive, respectively. Gray genes indicate false positives. **d**) Histograms representing the fraction of genes of different length classes relative to reference proteins, that were either predicted over at least 75%, 25-75%, or less than 25% of their expected length, or false positive predictions (FP). Protein sets and colors are the same as in a). Percentage labels in the plot indicate the total fraction per recall category for all length classes.

We found that on raw sequences less than 7% of the expected genes had good predictions, i.e. predictions that covered at least three-quarters of the expected gene length. 45% of the expected genes were missing entirely, and 48% of the predicted genes had very poor annotations covering less than a quarter of their expected length. Moreover, 12% of the predicted genes were false positives that had no match to the reference. By contrast, for frameshift corrected sequences, 84% of all expected genes were predicted, and 77% over three-quarters of their expected length. At the same time, the number of false-positive predictions dropped below 5%.

If broken down further by different gene-length categories, we found that short genes (<150 codons) were more often either missed entirely or annotated in full, while longer genes more often were present as fragments, and on raw sequences were never recovered at full-length. These differences are expected, as the disruption of an open reading frame is directly proportional to its length. In the given dataset we annotated 1,410 frameshifts in 40,656 aligned codons, which translates to an average of about 1 frameshift in 30 codons, which is very close to the minimum length required by most gene prediction programs. This suggests a hit-or-miss for many genes in the below 150 codon size range, consistent with our observations.

### Real bacterial nanopore long-reads corrected with different reference databases

To test our method on real data, and to also assess the effect of protein divergence, we turned to a second dataset - a recently published set of nanopore long reads obtained from the pathogenic bacterium *Mycoplasma bovis*^*33*^. The observed error rates were 1.6% substitutions, 1.5% insertions and 2.5% deletions. Analogous to the viral test set, we called genes on an *M. bovis* reference genome, as well as on raw and corrected reads. In this case, however, we tested three different protein sets for the correction: the best matching proteins from the UniRef90 database^35^, as well as the best matching UniRef90 proteins with at most 80% or at most 60% identity to any protein from the reference genome. Mean amino acid identities compared to the reference for these sets were 97%, 72%, and 58%, respectively (Fig. 2c/d).

The results we obtained were consistent with those from the simulated viral dataset. On raw reads only 2% of genes had good predictions, 36% of expected genes were missing, and 6% of the predicted genes were false positives. For the UniRef90 sets - best match, best match with less than 80% identity, and best match with less than 60% identity - the recall of largely complete genes was 74%, 66%, and 54%, respectively. 10%, 13% and 19% of genes were missed, and false-positive rates were 2%, 3% and 4%, respectively.

These results show that frameshift correction can improve gene prediction on long-read data considerably. However, we also note that similar to the simulated dataset, a fair proportion of primarily short genes could not be predicted, even if the correction was performed with highly similar reference proteins. We attribute this observation to two circumstances: 1) random errors in some short genes preventing significant nucleotide-to-protein alignments in the first place, thus, leaving the respective regions uncorrected, impairing gene calls; 2) obscured regulatory motifs outside coding regions, which, in particular, for short genes provide important additional evidence for the prediction program to distinguish spurious from truly coding open reading frames. Because our approach only uses guide proteins for correction, errors in untranslated regions can not be identified and can remain a problem for annotation.

Nevertheless, our results demonstrate that even with distantly related reference proteins, a significant proportion of the data can be corrected to a point where meaningful gene calls can be obtained.

### Metagenome-assembled genomes from closely related strains

We next investigated the impact of proovframe correction on long-read-assembled genomes by way of a gene-centric pangenome analysis. Since spurious amino acid sequences that emerge from frameshift errors have a substantial impact on comparative genomics analyses, pangenomics offers an ideal way to assess the efficacy of any correction algorithm. We focused on three complete and circular genomes of the Verrucomicrobium *Akkermansia*, which we have assembled from the long-read sequencing of three fecal samples collected from a single individual who received Fecal Microbiota Transplantation (FMT)^36^. Genome W0 originated from the pre-FMT sample, whereas genomes W1 and W48 originated from 1-week and 48-weeks after FMT. The previous analysis of these data concluded that *Akkermansia* W0 is replaced by *Akkermansia* W1 after FMT and persisted in the recipient for at least 48 weeks^36^. Indeed, the genomes W1 and W48 resolved to *Akkermansia muciniphila*, and the pairwise genome-wide average nucleotide identity (gANI) between them was over 99.9%, confirming that these genomes likely represent the same population. In contrast, the genome W0 shared a gANI score of only 88% with W1 and W48, and likely represents a novel species of *Akkermansia* as we failed to assign taxonomy to this genome below the genus level.

This collection of three genomes offers a powerful real-world test case: the high level of similarity between W1 and W48 suggests that the number of singleton gene clusters (genes unique to either one genome) should be minimal, if not zero, in any pangenomic analysis. Along the same lines, W0, W1, and W48 must share a relatively large number of genes since they are in the same genus. Finally, the absence of a reference genome for W0 gives a unique opportunity to test the efficacy of proovframe against relatively novel metagenome-assembled genomes with no representatives in public genome databases. We computed four pangenomes using the gene calls from the long-read assemblies (1) without further refinement, (2) after correction with proovframe, (3) after polishing with Illumina short-reads, and (4) after polishing with Illumina short-reads followed by proovframe correction.

The first pangenome from the unrefined long-read assembly revealed a substantial amount of random frameshift errors across all genomes where the number of gene clusters that represented single-copy core genes was small and the number of singleton gene clusters (in W1 and W48) was high relative to pangenomes with corrections (Fig. 3). Correcting the *Akkermansia* genomes either with proovframe or using Illumina short reads alone improved these metrics dramatically: the number of single-copy core genes increased from 1,488 to 1,695 and 1,642, and singleton gene clusters decreased from 1,593 to 177 and 38, respectively (Fig. 3). Across closely related populations, the alignment of gene amino acid sequences that are similar enough to be grouped together into the same gene cluster in a pangenome will typically lack any gap characters due to the constrained nature of the amino acid sequence space. However, fragmented genes or gene calls that suffer from partial frameshift errors can increase the number of gap characters observed in gene clusters dramatically. Indeed, anvi’o reported 3,595 gene clusters with a substantial number of gap characters for the pangenome with the unprocessed long-read assembly. Yet, correcting assembled genomes either with proovframe or using Illumina short reads alone reduced this number to 512 and 600, respectively (Fig. 3). These results show that the quality improvements of proovframe are comparable to polishing long-read assemblies with Illumina short reads. Our results also showed that combining proovframe correction and short-read polishing offered the best results, where the singleton gene clusters in W1 and W40 reduced from 1,593 to 15, the number of gene clusters with substantial heterogeneity reduced from 3,595 to 153, and the number of single-copy core genes between the three genomes increased from 1,488 to 1,916. Overall, these results suggest that proovframe substantially improves insights from pangenomes of genomes reconstructed from complex metagenomes both with and without Illumina short reads.

**Figure 3.**
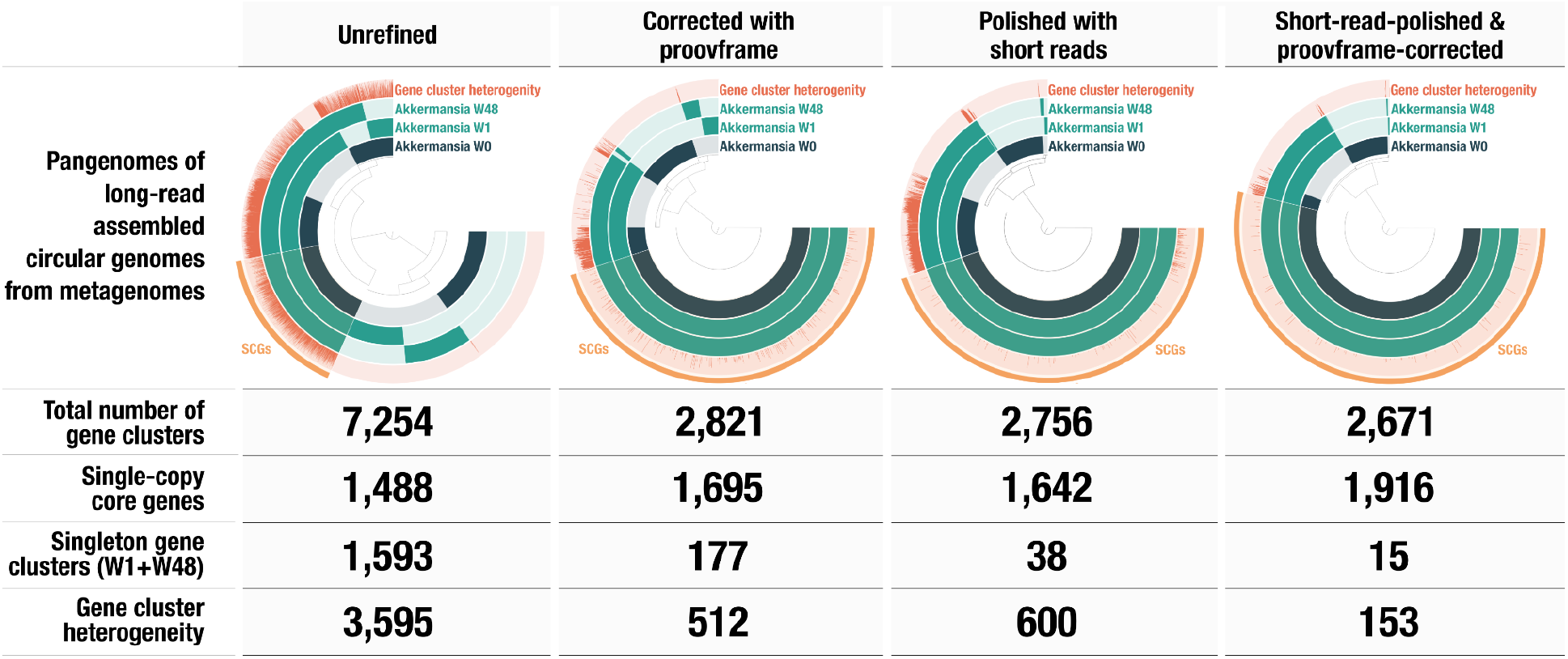
A set of *Akkermansia* pangenomes with and without short-read polishing and proovframe correction. Each column represents the analysis of the same set of circular *Akkermansia* genomes assembled from MinION long-reads. The first row displays the pangenomes, where the three concentric inner layers correspond to *Akkermansia* genomes where each item organized by the dendrograms in the center of each pangenome represents gene clusters. The intensity of the color indicates the presence of a given gene cluster in each genome. The ‘gene cluster heterogeneity’ layer quantifies the proportion of gap characters in the aligned amino acid sequences in a given gene cluster to the number of amino acid sequence residues. High heterogeneity indicates poor gene alignment with a high number of gaps, and a score of 0 indicates no gaps in the alignment. For each pangenome the table below reports the number of single-copy core genes, the number of singleton gene clusters for *Akkermansia* W1 and W48, and the number of gene clusters with a heterogeneity score above 0.1.

### Nanopore long reads from a highly diverse marine small-particle-fraction metagenome

Finally, to test the applicability of frameshift correction to complex environmental metagenomic data, we applied it to a highly diverse set of nanopore long reads we had recently obtained from a marine small-particle-fraction metagenomic sample containing primarily extracellular vesicles and tailless viruses. Pairwise read-to-read similarities indicated that fewer than 5% of the raw reads had sufficient coverage for assembly and consensus polishing, and thus an alternative correction strategy was essential for a comprehensive analysis.

For the correction, we used a protein database that we had assembled with the protein assembler PLASS^46^ from an accompanying Illumina HiSeq library. As before, we identified reads representing viruses with virsorter2^47^ on both the raw and the corrected reads and annotated the reads flagged as viral with Prodigal^31^. With frameshift-corrected reads we were able to identify 1.8 times as many viral reads, carrying twice as many genes with a 2.5 times greater average gene length. The coding density for corrected reads was 92%, while it was only 54% on raw reads. As viral genomes typically have coding densities of 90% or more, our results suggest that we successfully recovered the majority of the genes encoded on these putatively viral sequences.

### Remarks on runtime

Proovframe’s runtime is primarily determined by the initial DIAMOND alignment step. Aligning 120 M proteins from UniRef90 with DIAMOND in default ‘fast’ mode on a 60-core compute node took 20 minutes for the seven different *Akkermansia* genomes. Aligning 43 M proteins from the marine metaproteome took 4h for 3.3 Gbp of our small-particle-fraction nanopore dataset, and 10 hours for the same read set in the ‘more-sensitive’ mode. The subsequent correction of the reads took a few seconds and a few minutes, respectively. We did not perform any more precise or comprehensive runtime benchmarks because we expect runtimes for different projects to be highly variable depending on the specific experimental design and scientific question. Information on expected runtimes, in general, can be extrapolated from DIAMOND’s performance on the different query and target databases, sensitivity modes, and data types investigated in other studies^23,48^. Our small-particle-fraction dataset likely marks an upper boundary for a realistic proovframe use-case because we used a very large protein database, tested a sensitive alignment mode, and our reads comprised more than 90% coding regions. Therefore, we expect that the majority of projects can be processed with proovframe on similar state-of-the-art computing infrastructures within a reasonable timeframe of hours to days.

## Conclusion

*Proovframe* efficiently corrects frameshift errors in long-read sequencing reads and derived long-read assemblies. It considerably improves open reading frame fidelity and, in turn, gene prediction and overall analyzability of long-read data. In contrast to typical polishing strategies, which rely on sufficient sequencing depth in either the long-reads themselves or accompanying Illumina short-read data, proovframe utilizes alignments to reference proteins as guides for the correction. It can, thus, be applied at no additional costs other than computing time to a wide array of long-read sequencing projects, including complex samples, such as environmental microbiomes, where polishing is not or at least only in part an option.

The main limitation of the protein-guided correction approach compared to classic polishing is that the restoration of frame fidelity has only little effect on the actual per-base nucleotide level accuracy of the underlying data: Proovframe only operates on aligned coding regions, does not correct substitution errors, and only inserts unspecific nucleotides (N/NN for deletions, NNN for premature stops). Our main objective is the conservative restoration of open reading frames to facilitate gene prediction and gene-based downstream analysis. In some cases, this also means that proovframe can mask genuine frameshifts, for example, in pseudogenized genes. Moreover, proovframe requires an adequate reference protein set. For most projects, however, this should not be too limiting given that the correction also works well with remote protein homologs (<60% identity for the *Mycoplasma* benchmark set, Fig. 2) and even comprehensive databases such as UniRef90, which can be aligned within reasonable timeframes.

On long-read assemblies, as demonstrated for the metagenome-assembled *Akkermansia* pangenomes, proovframe can perform comparably to polishing with Illumina short reads (Fig. 3). Moreover, our analyses showed that combining polishing and proovframe resulted in the most substantial improvement of the assessed metrics, thus indicating that running proovframe as an extra step for already polished data could be a generally advisable practice.

In contrast to other methods, proovframe can also be used directly on raw reads, thus enabling the analysis of individual reads and even entire samples which otherwise would not be accessible to standard gene-based downstream analyses: Applied to a small-particle-fraction long-read metagenome with many reads representing full-length but unique viruses, we identified almost five times more viral proteins sequences after the correction, providing us with a much more complete picture of this complex virome.

In summary, proovframe provides a simple and efficient way to overcome frameshift errors in long-read data, both as an alternative or as a complementary approach to long- and short-read polishing. We expect that it can be applied to a wide variety of projects and, thus, will help to accelerate long-read-based genomics research across different fields and scales.

## Data and code availability

The datasets together with reproducible code for the benchmarking are available from GitHub (thackl/proovframe-benchmark) and Zenodo (10.5281/zenodo.5164669)^29^.

## Acknowledgments

TH and MGF acknowledge support from the Max Planck Society. A Helmsley Foundation grant to AME supported FT. AME acknowledges support from the Simons Foundation (#687269). EFD, JME, EL and AB were supported by grants from the Simons Foundation (grant #329108 and #721223), and the Gordon and Betty Moore Foundation (#3777). We thank I. Schlichting and C. Roome for support.

## Ethics declarations

The authors declare no competing interests.

## Author contribution

TH conceived the study, implemented the software and evaluated its general performance. FT and AME conceived and carried out the pangenome-based evaluation for the *Akkermansia* genomes. SJB, EFD, JME, EL, and AB conceived and implemented the development, collection, and production of the small-particle-fraction metagenomics long read and short read sequence data. TH wrote the manuscript with contributions from all authors. MGF supervised the project.

## References

1. Pollard, M. O., Gurdasani, D., Mentzer, A. J., Porter, T. & Sandhu, M. S. Long reads: their purpose and place. Hum. Mol. Genet. 27, R234–R241 (2018).

2. Fu, S., Wang, A. & Au, K. F. A comparative evaluation of hybrid error correction methods for error-prone long reads. Genome Biol. 20, 26 (2019).

3. Dohm, J. C., Peters, P., Stralis-Pavese, N. & Himmelbauer, H. Benchmarking of long-read correction methods. NAR Genomics and Bioinformatics 2, (2020).

4. Liu, X., Mei, W., Soltis, P. S., Soltis, D. E. & Barbazuk, W. B. Detecting alternatively spliced transcript isoforms from single-molecule long-read sequences without a reference genome. Mol. Ecol. Resour. 17, 1243–1256 (2017).

5. Ruan, J. & Li, H. Fast and accurate long-read assembly with wtdbg2. Nat. Methods 17, 155–158 (2020).

6. Kolmogorov, M. et al. metaFlye: scalable long-read metagenome assembly using repeat graphs. Nat. Methods 17, 1103–1110 (2020).

7. Quick, J. et al. Real-time, portable genome sequencing for Ebola surveillance. Nature 530, 228–232 (2016).

8. Rooke, S. Resolving complex mobile genetic elements with nanopore sequencing. Access Microbiology 1, (2019).

9. Nowoshilow, S. et al. The axolotl genome and the evolution of key tissue formation regulators. Nature 554, 50–55 (2018).

10. Palfalvi, G. et al. Genomes of the Venus Flytrap and Close Relatives Unveil the Roots of Plant Carnivory. Curr. Biol. 30, 2312–2320.e5 (2020).

11. Hackl, T., Duponchel, S., Barenhoff, K. & Weinmann, A. Endogenous virophages populate the genomes of a marine heterotrophic flagellate. bioRxiv (2020).

12. Slaby, B. M., Hackl, T., Horn, H., Bayer, K. & Hentschel, U. Metagenomic binning of a marine sponge microbiome reveals unity in defense but metabolic specialization. ISME J. (2017) doi:10.1038/ismej.2017.101.

13. McKenzie, S. K., Walston, R. F. & Allen, J. L. Complete, high-quality genomes from long-read metagenomic sequencing of two wolf lichen thalli reveals enigmatic genome architecture. Genomics 112, 3150–3156 (2020).

14. Beaulaurier, J. et al. Assembly-free single-molecule sequencing recovers complete virus genomes from natural microbial communities. Genome Res. 30, 437–446 (2020).

15. Chin, C.-S. et al. Nonhybrid, finished microbial genome assemblies from long-read SMRT sequencing data. Nat. Methods 1–9 (2013) doi:10.1038/nmeth.2474.

16. Hackl, T., Hedrich, R., Schultz, J. & Förster, F. proovread: large-scale high-accuracy PacBio correction through iterative short read consensus. Bioinformatics 30, 3004–3011 (2014).

17. Xiao, C.-L. et al. MECAT: fast mapping, error correction, and de novo assembly for single-molecule sequencing reads. Nat. Methods 14, 1072–1074 (2017).

18. Vaser, R., Sović, I., Nagarajan, N. & Šikić, M. Fast and accurate de novo genome assembly from long uncorrected reads. Genome Res. 27, 737–746 (2017).

19. Li, H. Minimap2: pairwise alignment for nucleotide sequences. Bioinformatics 34, 3094–3100 (2018).

20. Kolmogorov, M., Yuan, J., Lin, Y. & Pevzner, P. A. Assembly of long, error-prone reads using repeat graphs. Nat. Biotechnol. 37, 540–546 (2019).

21. Arumugam, K. et al. Annotated bacterial chromosomes from frame-shift-corrected long-read metagenomic data. Microbiome 7, 61 (2019).

22. Huson, D. H. et al. MEGAN-LR: new algorithms allow accurate binning and easy interactive exploration of metagenomic long reads and contigs. Biol. Direct 13, 6 (2018).

23. Buchfink, B., Xie, C. & Huson, D. H. Fast and sensitive protein alignment using DIAMOND. Nat. Methods 12, 59–60 (2015).

24. Haro-Moreno, J. M., López-Pérez, M. & Rodríguez-Valera, F. Long read metagenomics, the next step? Cold Spring Harbor Laboratory 2020.11.11.378109 (2020) doi:10.1101/2020.11.11.378109.

25. Chen, L.-X., Anantharaman, K., Shaiber, A., Eren, A. M. & Banfield, J. F. Accurate and complete genomes from metagenomes. Genome Res. 30, 315–333 (2020).

26. Fuhrman, J. A. Microbial community structure and its functional implications. Nature 459, 193–199 (2009).

27. Hackl, T. et al. Novel Integrative Elements and Genomic Plasticity in Ocean Ecosystems. Cell preprint (2021) doi:10.2139/ssrn.3817805.

28. Bernheim, A. & Sorek, R. The pan-immune system of bacteria: antiviral defence as a community resource. Nat. Rev. Microbiol. 18, 113–119 (2020).

29. Hackl, T. thackl/proovframe-benchmark: proovframe-benchmark-v3.0. (2021). doi:10.5281/zenodo.5164669.

30. Yang, C., Chu, J., Warren, R. L. & Birol, I. NanoSim: nanopore sequence read simulator based on statistical characterization. Gigascience 6, 1–6 (2017).

31. Hyatt, D. et al. Prodigal: prokaryotic gene recognition and translation initiation site identification. BMC Bioinformatics 11, 119 (2010).

32. Hackl, T. & Ankenbrand, M. J. gggenomes - A grammar of graphics for comparative genomics. (2021).

33. Vereecke, N. et al. High quality genome assemblies of Mycoplasma bovis using a taxon-specific Bonito basecaller for MinION and Flongle long-read nanopore sequencing. BMC Bioinformatics 21, 517 (2020).

34. Silvestre-Ryan, J. & Holmes, I. Pair consensus decoding improves accuracy of neural network basecallers for nanopore sequencing. Genome Biol. 22, 38 (2021).

35. Suzek, B. E. et al. UniRef clusters: a comprehensive and scalable alternative for improving sequence similarity searches. Bioinformatics 31, 926–932 (2015).

36. Watson, A. R. et al. Adaptive ecological processes and metabolic independence drive microbial colonization and resilience in the human gut. bioRxiv 2021.03.02.433653 (2021) doi:10.1101/2021.03.02.433653.

37. Trigodet, F. et al. High molecular weight DNA extraction strategies for long-read sequencing of complex metagenomes. bioRxiv 2021.03.03.433801 (2021) doi:10.1101/2021.03.03.433801.

38. Walker, B. J. et al. Pilon: an integrated tool for comprehensive microbial variant detection and genome assembly improvement. PLoS One 9, e112963 (2014).

39. Delmont, T. O. & Eren, A. M. Linking pangenomes and metagenomes: the Prochlorococcus metapangenome. PeerJ 6, e4320 (2018).

40. Eren, A. M. et al. Community-led, integrated, reproducible multi-omics with anvi’o. Nat Microbiol 6, 3–6 (2021).

41. Buchfink, B., Reuter, K. & Drost, H.-G. Sensitive protein alignments at tree-of-life scale using DIAMOND. Nat. Methods 18, 366–368 (2021).

42. van Dongen, S. & Abreu-Goodger, C. Using MCL to Extract Clusters from Networks. in Bacterial Molecular Networks: Methods and Protocols (eds. van Helden, J., Toussaint, A. & Thieffry, D.) 281–295 (Springer New York, 2012). doi:10.1007/978-1-61779-361-5_15.

43. Edgar, R. C. MUSCLE: a multiple sequence alignment method with reduced time and space complexity. BMC Bioinformatics 5, 113 (2004).

44. Biller, S. J. et al. Bacterial vesicles in marine ecosystems. Science 343, 183–186 (2014).

45. Shen, W., Le, S., Li, Y. & Hu, F. SeqKit: A Cross-Platform and Ultrafast Toolkit for FASTA/Q File Manipulation. PLoS One 11, e0163962 (2016).

46. Steinegger, M., Mirdita, M. & Söding, J. Protein-level assembly increases protein sequence recovery from metagenomic samples manyfold. Nat. Methods 16, 603–606 (2019).

47. Roux, S., Enault, F., Hurwitz, B. L. & Sullivan, M. B. VirSorter: mining viral signal from microbial genomic data. PeerJ 3, e985 (2015).

48. Hernández-Salmerón, J. E. & Moreno-Hagelsieb, G. Progress in quickly finding orthologs as reciprocal best hits: comparing blast, last, diamond and MMseqs2. BMC Genomics vol. 21 (2020).

